# Multitrophic assembly: a perspective from modern coexistence theory

**DOI:** 10.1101/2023.03.20.533409

**Authors:** Chuliang Song, Jurg W. Spaak

## Abstract

Ecological communities encompass rich diversity across multiple trophics. While modern coexistence theory has been useful in understanding community assembly, its traditional formalism only allows for the study of assembly within a single trophic level. Here, using an expanded definition of niche and fitness differences applicable to multi-trophic communities, we study how diversity within and across trophics affect species coexistence. Specifically, we investigate how assembly in one trophic level impacts the coexistence of three types of communities: (1) the single-trophic subcommunity with species at that level, (2) the single-trophic subcommunity with species at an adjacent level, and (3) the entire multitrophic community. We find that while coexistence mechanisms are similar for single-trophic communities, they differ for multitrophic ones. We also find that fitness differences primarily constrain diversity in lower-level tropics, while niche differences primarily constrain diversity in higher-level tropics. Empirical data corroborates our predictions about multitrophic structures. Our work provides needed theoretical expectation of multitrophic communities within modern coexistence theory.

## Introduction

A fundamental characteristic of eco-complexity is that species interact within and across trophic levels (Beckage *et al*., 2011; Godoy *et al*., 2018). Ample empirical evidence suggests that these multitrophic structures strongly affect the patterns of community assembly (Drake, 1991; Price & Morin, 2004; Olito & Fukami, 2009; Song *et al*., 2018a; Pringle *et al*., 2019). Thus, understanding how these multitrophic structures regulate community assembly is a central question in community ecology with direct implications for conservation and restoration of natural ecosystems (Wratten *et al*., 2000; Gossner *et al*., 2016; Eisenhauer *et al*., 2019).

Yet, our current understanding of community assembly has been mostly shaped by (often implicit) separation of trophic levels (Figure 1; Seibold *et al*. 2018). To understand community assembly, the most frequent approach focuses on how a new invading species affects its competitors in the same trophic level (Chase & Leibold, 2003; Letten *et al*., 2017; Shoemaker *et al*., 2020). We denote this the *traditional focus* (Fig. 1). In contrast, another approach focuses on how this invader affects other species on the higher trophic level, or more generally an adjacent trophic level (Petry *et al*., 2018; Terry *et al*., 2021). We denote this the *alternative focus* (Fig. 1). However, such separation of trophic levels is not always justified (Levine *et al*., 2017; Godoy *et al*., 2018; Spaak *et al*., 2021c). The inclusion of multiple trophic levels strongly affects our view of community assembly. For example, if an invading species excludes another species from the same or different trophic level, this exclusion can further affect species throughout the entire network, known as secondary extinctions (Brodie *et al*., 2014). In addition, many ecological properties can only be studied for the community as a whole, such as link-species relationships (Carpentier *et al*., 2021), distribution of biomass across trophic levels (Galiana *et al*., 2021), or average food-chain length (Post, 2002). Thus, the focus on the multi-trophic community as a whole is the most relevant scale for many ecological questions (Fig. 1).

**Figure 1:**
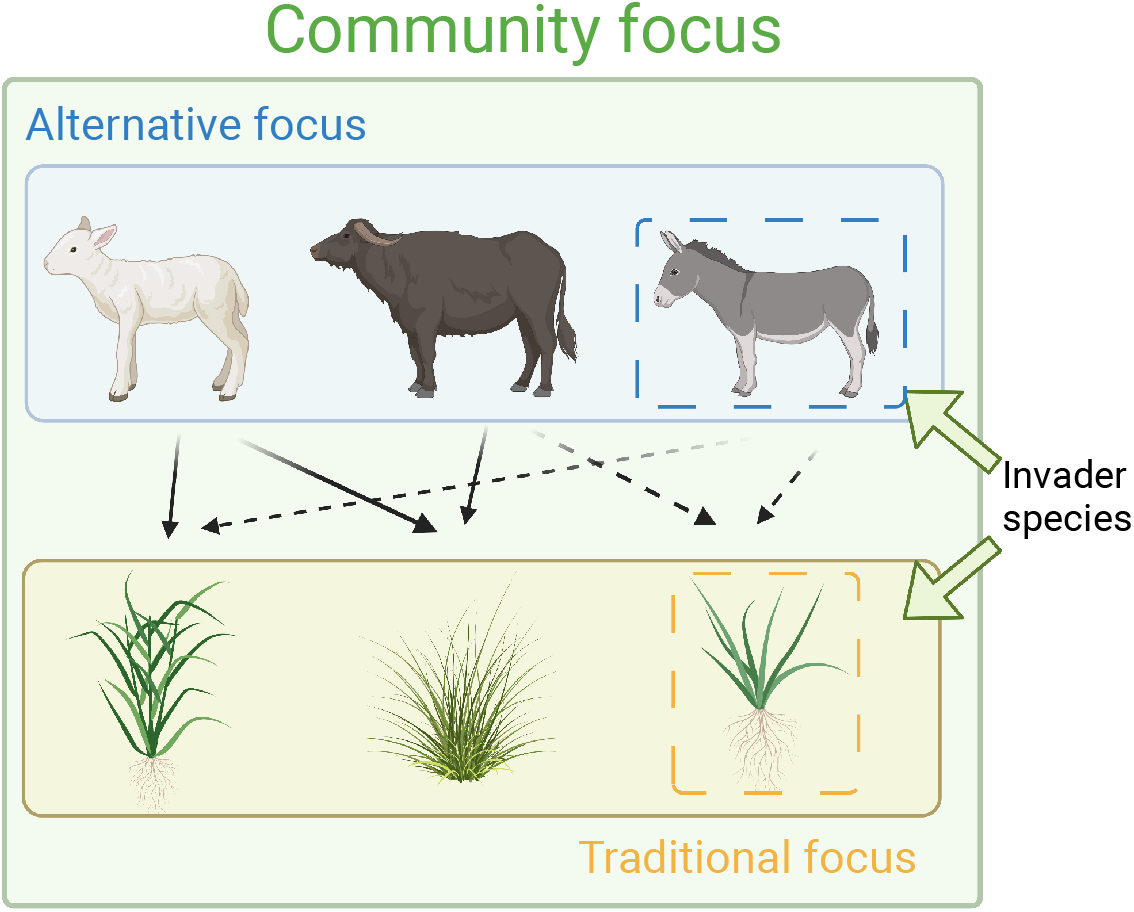
Different perspectives on multitrophic assembly. We consider a hypothetical, two-trophic community with 3 basal species and 3 consumer species. Traditionally, we study the impact of an invader or increased species richness on the invaded trophic level (Traditional focus). This effect is usually negative due to heightened competition for resources or predator pressure. Alternatively, we may be interested in how an invader affects an adjacent trophic level (Alternative focus), which generally has a positive effect by increasing resource availability. However, it is essential to investigate the impact of invaders on the entire community (Community focus) since their effects can be both positive and negative.

Despite an emergent line of theoretical frameworks on multitrophic structures (Barabás *et al*., 2016; Song *et al*., 2018b; Wang & Brose, 2018; Koffel *et al*., 2021), modern coexistence theory—a key theoretical framework widely adopted by empiricists—has been an exception. The majority of empirical studies using modern coexistence theory has primarily focused on single trophic level with competition (reviewed in Barabás *et al*. 2018). This problem is partly due to the limitations in the theoretical framework: niche and fitness differences—fundamental concepts in modern coexistence theory—are not well-defined for multitrophic systems until recently (Spaak *et al*., 2022b). Specifically, in the canonical formalism of modern coexistence theory, niche differences measure the overlap in resource use between two competitors, while fitness differences measured the difference in total resource consumption as well as differences in mortality (Chesson, 1990; Chesson & Kuang, 2008). Recently, Spaak *et al*. (2021c) generalized niche and fitness differences that are applicable multitrophic communities. In their definition, niche differences measure how similar the interspecific competition is to intraspecific competition, while fitness differences measure how well a species would do if all other species occupied exactly the same niche as the focal species. These new measures of niche and fitness differences serve as a common currency to compare coexistence mechanisms across different trophic levels.

Here, we apply modern coexistence theory to establish a null expectation of how multitrophic structures modulate ecological assembly across trophic levels. Our study focused on two questions: (1) How does a change in community composition affect the coexistence of the community? and (2) Are the mechanisms driving coexistence in different trophic levels the same? To answer these questions, we first computed niche and fitness differences, both by analytic mathematics and simulations, of two-trophic communities with varying community richness and percentage of basal species. We analyzed how an invasion of a species affects niche and fitness differences at different communities: the subcommunity with species at the same level only (traditional focus), the subcommunity with species at adjacent trophic level only (alternative focus), and the whole multitrophic community *community focus). Analyzing all these communities suggest that we should expect a balanced food web due to an inevitable tradeoff: community composition affects niche differences in higher trophic levels, while it affects fitness differences in the basal trophic level. However, the traditional and alternative focus find that coexistence mechanism do not differ between trophic level, while the community focus does differ. We then generalized our analysis by extending to three-trophic communities. Finally, we tested our predictions using empirical multi-trophic network data.

## Methods

### Community model

We consider communities with two trophic levels. We assume that community dynamics is governed by the Lotka-Volterra model

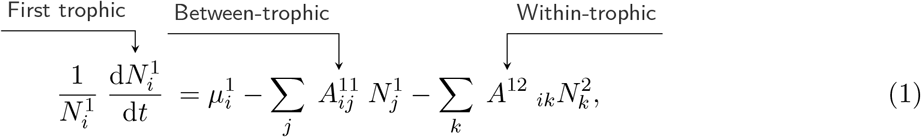

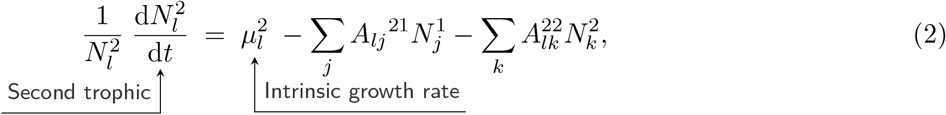

where 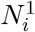 is the density of species *i* in the first trophic level, 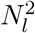 is the density of species *l* from the second trophic level, 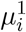 and 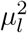 denote the intrinsic growth rates of species in the first and second trophic levels. Similarly, *A*^11^, *A*^12^, *A*^21^ and *A*^22^ denote the interaction matrices between and within the respective trophic levels. *A*^11^ and *A*^22^ capture all interactions with other, not explicitly mentioned, trophic levels, such as resources for *A*^11^ (MacArthur, 1970) or higher trophic levels for *A*^22^ (Chesson & Kuang, 2008). More generally, *A*^11^ and *A*^22^ can capture all non-trophic interactions such as competition for space (Shoemaker *et al*., 2019), breeding opportunities, and other direct species interactions (Kawatsu *et al*., 2021). For symmetry, we choose to have minus signs for all species interactions *A^xy^* and positive signs in front of intrinsic growth rates *μ^x^*.

For the simulations and the analytical results we take a generally adopted simplifying assumption that the community assembles from a random species pool (Allesina & Tang, 2015; Bunin, 2017; Serván *et al*., 2018). Specifically, we assume that all interaction strengths are chosen at random and independent of other species. We also assume that the average strength of the interaction does not depend on the trophic level, i.e. 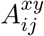 are all drawn from the same underlying distribution. This assumption is likely wrong, however it enables us to understand how the trophic structure affects community stability, as opposed to differences in interaction strength. These entries are all drawn independent of each other, except for the correlation of predation and consumption. We assume that consumption 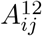 and predation 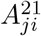 are identical (i.e., *A*^21^ = −(*A*^12^)^*T*^). Finally, we assume that interspecific competition is 1 for all species, i.e. 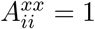.

### Niche and fitness differences

Niche and fitness differences have multiple definitions under the umbrella of modern coexistence theory (Chesson, 2003; Adler *et al*., 2007; Carroll *et al*., 2011; Godoy *et al*., 2014; Zhao *et al*., 2016; Saavedra *et al*., 2017; Carmel *et al*., 2017; Bimler *et al*., 2018; Spaak & De Laender, 2020). Here, we adopt the definition proposed by Spaak & De Laender (2020) with adjustments made by Spaak *et al*. (2021c). This definition is currently the only one out of all the definitions that can operate on multi-species and multi-trophic communities which agrees with intuitive undersanding of faciliation or competition (Spaak *et al*., 2021d). While this definition works for more general population dynamics, we limit our focus on the Lotka-Volterra community model (Eqns. 1 and 2). The computation of niche and fitness differences is based on *invasion growth rates*, the species *i* growth rate while invading the resident community at equilibrium. We denote the equilibrium *N*^(−*i*,*)^ = (*A*^−*i*,−*i*^)^−1^*μ*^−*i*^, where *A*^−*i*,−*i*^ and *μ*^−*i*^ are the interaction matrix and the intrinsic growth rates with row and/or column *i* removed. With this notation and Lotka-Volterra community model, the invasion growth rate *r_i_* is given by 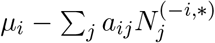. To normalize niche and fitness differences, the invasion growth rate *r_i_* is compared to two hypothetical invasion growth rates. The first one is the intrinsic growth rate *μ_i_*, which can be interpreted as the invasion growth rate if species *i* did not interact with any other species (i.e., *a_ij_* = 0 for all *j*). The second one is the no-niche growth rate *η_i_*, which is the invasion growth rate if all species had the same niche as species *i*. However, changing *a_ij_* to *a_ii_* does not only change the niche of species *j*, but also the fitness of species *j* (Chu & Adler, 2015). The noniche growth rate *η_i_* is therefore the growth rate if we set 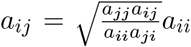, where 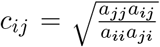 is the conversion factor converting species *i* to species *j* (see Spaak & De Laender 2021 for a detailed derivation).

Given these three invasion growth rates (*μ_i_*,*r_i_* and *η_i_*), the niche difference 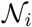 and fitness difference 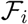 are defined by

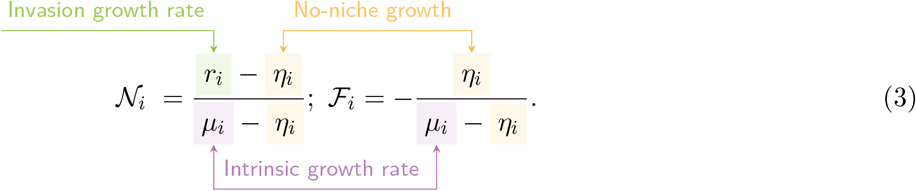

To see why these definitions are ecologically intuitive, we can take a look at the following cases. Focusing on niche difference, 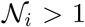 implies that species benefit from the presence of other species (i.e. *μ_i_* < *r_i_* Spaak & De Laender 2020), while 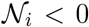 implies species have stronger interspecific than intraspecific interactions (i.e. *η_i_* > *r_i_*; Ke & Letten 2018). Then focusing on fitness difference, 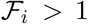 implies that species depend on other species such as predation interactions (i.e. *μ_i_* < 0; Spaak *et al*. 2021c), while 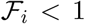 implies that a species can grow in the absence of other species (i.e. *μ_i_* > 0; Spaak *et al*. 2021c). Finally, a species has a positive invasion growth rate if its niche difference overcomes its fitness difference, i.e. 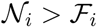 (Adler *et al*., 2007; Chesson, 2000).

### Perspectives on multitrophic communities

We analyzed how community assembly affects stability by examining a community under three different focuses: The traditional focus considers the trophic level in which a new species invades, the alternative focus looks at the adjacent trophic level, and the community focus examines all trophic levels simultaneously.

To calculate niche and fitness differences for the entire community, we can directly apply methods of niche and fitness differences to the community model (Spaak *et al*., 2021c). However, if we want to concentrate on a specific trophic level, we treat species from other levels as limiting factors. As niche and fitness differences are calculated from three growth rates evaluated at steady states, we can therefore solve these equations by setting the growth rates of the limiting factors to 0. This approach is mathematically equivalent to using timescale separation from MacArthur resource model (MacArthur, 1970; O’Dwyer, 2018). A key ecological difference, though, is that we do not assume that different trophic levels have different intrinsic time scales of ecological processes, rather we only compute growth rates after a steady state is reached. Mathematically, this process is equivalent to studying the effective Lotka-Volterra dynamics (Appendix A):

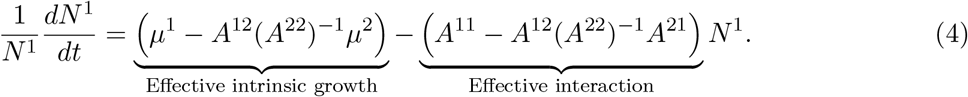

The methods discussed in Spaak *et al*. (2021c) can then be applied to this altered community model.

We analytically computed niche and fitness differences for the traditional and alternative focus under the assumption that all interspecific species interactions are identical. Additionally, we computed how the community focus depends on changes in species richness in the lower or higher trophic level.

### Simulations

To relax the simplifying assumptions in analytic arguments, we explore more realistic structures using simulations. Our focus was on how multitrophic structure affects coexistence of different trophic levels, so we removed variations within a trophic level that are not intrinsically linked to their trophic level. Specifically, we assumed that all interspecific interactions are equally strong and only differ in their sign (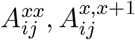 and 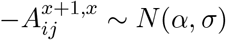) independent of the trophic level *x*. All intraspecific interactions are drawn from different distributions 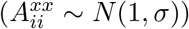, again independent of the trophic level *x*. Similarly, the intrinsic growth rate of all non-basal species were drawn from the same underlying distribution, 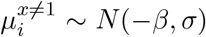 However, the intrinsic growth rate of the basal species stems from a different distribution, because positive intrinsic growth is intrinsically linked to its trophic position 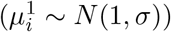.

We then test whether our findings in two-trophic communities generalize to tri-trophic community. Specifically, for a community with 21 species in total, we altered the proportion of each trophic level and computed niche and fitness differences of the community as a whole (community focus). We then computed for each species *i* how strongly the niche differences compensates for the fitness differences 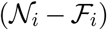 and defined the stability of a community as that of its least stable species (min 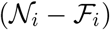).

Finally, to explore the most stable multitrophic structures, we randomly chose a mean interaction strength *α*, standard-deviation of interaction strength *σ*, species richness *n* and mortality rate of higher trophic level *m_i_*. For each random parametrization, we kept the five most stable community compositions.

### Empirical data

To confront our theoretical results with empirical data, we analyzed 358 food-webs from web of life (WWW.WEB-OF-LIFE.ES). We determined the trophic level of each species. We first considered species without prey as basal and assigning them a trophic level of 1. For all other species, we calculated their trophic level by taking the mean of their prey’s trophic levels and adding 1. We grouped species with trophic levels between 2 and 2.5 into the second trophic level, while those exceeding 2.5 were placed in the third trophic level. We then compared the distribution of the community composition of empirical trophic networks with theoretical predictions from three-trophic communities.

## Results

### Assembly in two-trophic communities

We first examine how assembly in one trophic level affects stability within that same level (traditional focus in Figure 1). Higher species richness within a trophic level increases fitness differences among species (Figure 2 D,J), but does not affect their niche differences (Panels A,G). This pattern holds across all trophic levels (Appendix B) and generalizes earlier findings by Spaak *et al*. (2021a) for multi-trophic communities. A heuristic explanation is that fitness differences reflect the competitive strength of a focal species relative to its competitors, and increase with the number of competitors. On the other hand, niche differences compare the niche of a focal species to the average niche of its competitors and are therefore unaffected by species richness (Spaak *et al*., 2021a).

**Figure 2:**
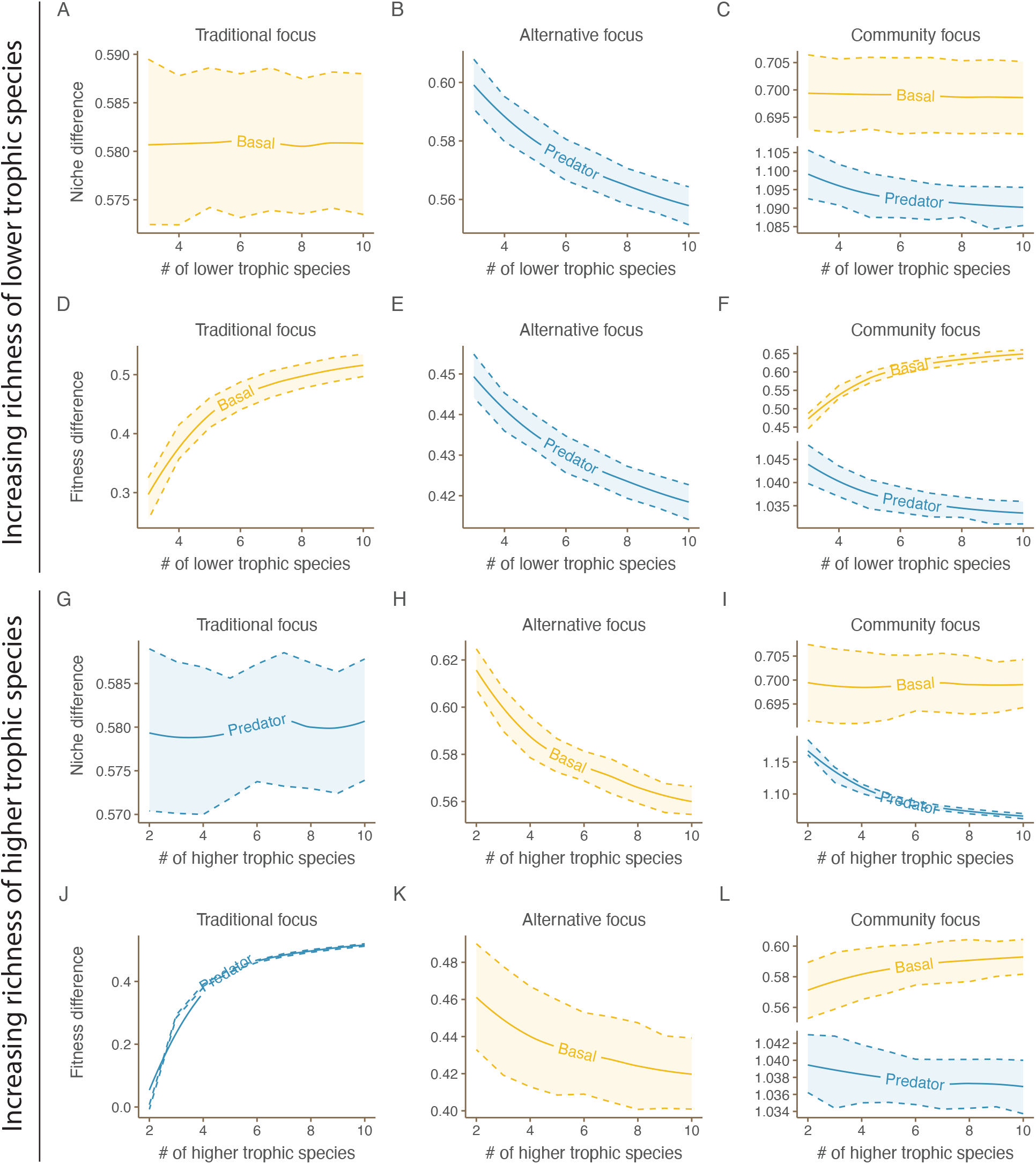
How species richness affects niche and fitness differences. The first two rows show a change in basal species richness, while the last two rows show a change in predator species richness. According to the traditional focus (the first column), increasing species richness does not affect niche differences, but increases fitness differences. According to the alternative focus (the second column), increasing species richness decreases both niche and fitness differences, whereas the effect on niche differences is slightly stronger. Finally, according to the community focus (the third column), niche differences of the basal species are not affected, while fitness differences of the basal species increase. Both niche and fitness differences of the predator species decrease with increasing species richness. Note that y-axes have drastically different ranges, indicating different magnitudes of the effects.

We then examine how assembly in one trophic level affects the stability of the adjacent trophic level (alternative focus in Figure 1). Increasing species richness in a trophic level reduces niche differences (Fig. 2B & H) and fitness differences (Fig. 2E & K) of the adjacent trophic level. A heuristic explanation behind this pattern is that the effective interaction between species in the focal trophic level is a combination of actual inter-trophic interactions *A*^11^ and interactions stemming from (apparent) competition (*A*^12^ (*A*^22^)^−1^ *A*^21^). Our parametrization assumes that species in the focal trophic level have on average identical interactions with all species from the adjacent trophic level, making the (apparent) competition term for these species neutral. Moreover, increasing species richness in the adjacent level increases the strength of the (apparent) competition, making it more neutral and decreasing niche and fitness differences. More careful mathematical analysis shows that increasing species richness has a stronger effect on niche than on fitness differences; specifically, rate of change of fitness differences is (1 − 1/*n*_1_) times that of niche differences (see Appendix C), where *n_1_* represents number of species at basal tropic levels. This difference is most pronounced when communities have few species.

Lastly, we analyze the community-level effects of assembly on coexistence by computing niche and fitness differences for all species in a community simultaneously (community focus in Figure 1). We first focus on basal species. We find that increasing species richness, either of the higher or lower trophic level, does not affect niche differences of the basal species (Fig. 2C & I), but increases their fitness differences (Fig. 2F & L and Appendix D). Intuitively, niche differences are a weighted average of the pair-wise niche differences a species has with the other species (Spaak *et al*., 2021b), which is not affected by species richness for the basal species. In contrast, fitness differences species richness is the weighted sum of the pair-wise fitness differences (Spaak *et al*., 2021d), which increases with increasing species richness as the sum has more terms. We then focus on higher trophic species. We find that increasing species richness decreases both niche (Fig. 2C & I) and fitness differences (Fig. 2F & L) of the higher trophic level. Intuitively, increasing species richness of the higher trophic level decreases the total biomass of the lower trophic level due to increased predation, but increases the total biomass of the higher trophic level due to overyielding. Therefore, increasing species richness in the higher trophic level decreases niche differences of the higher trophic level, as more interspecific interactions are competitive (low niche difference) and fewer interactions are predation (high niche differences). Interestingly, increasing the species richness in the lower trophic level also decreases niche differences of the higher trophic level. While higher species richness of the lower trophic level increases the biomass of the lower trophic level due to overyielding it even more strongly increases the total biomass of the higher trophic level due to predation. Again, a random interaction with another species is more likely to be competitive than to be a predator prey interaction. The fitness differences are driven by total biomass, which increases with both increasing species richness of the lower and of the higher trophic level. However, for the higher trophic level this implies a decrease in species richness, as fitness differences exceed 1.

### Balanced structure in two-trophic communities

Here, we study how community composition affects niche and fitness differences by manipulating the proportion of basal species while keeping total species richness constant. We did not use traditional or alternative focus here because altering the proportion of basal species affects both higher and lower trophic levels, resulting in no changes to overall species richness.

Focusing on niche differences, increasing the proportion of basal species does not affect the niche differences of lower trophic levels, but increases the niche differences of higher trophic levels (Fig. 3A) Heuristically, niche difference is calculated as the mean difference between a species and a randomly chosen individual from the community. A predator species with a higher proportion of basal species will interact more frequently with prey, leading to an increase in its niche differences. However, in our simulations, all other species have the same niche difference with a basal species regardless of their trophic level (see Methods), so the proportion of basal species does not affect its niche difference. The effect on basal species’ niche differences depends on whether their differences with predators are smaller or larger than those with another basal species; this effect is generally small.

**Figure 3:**
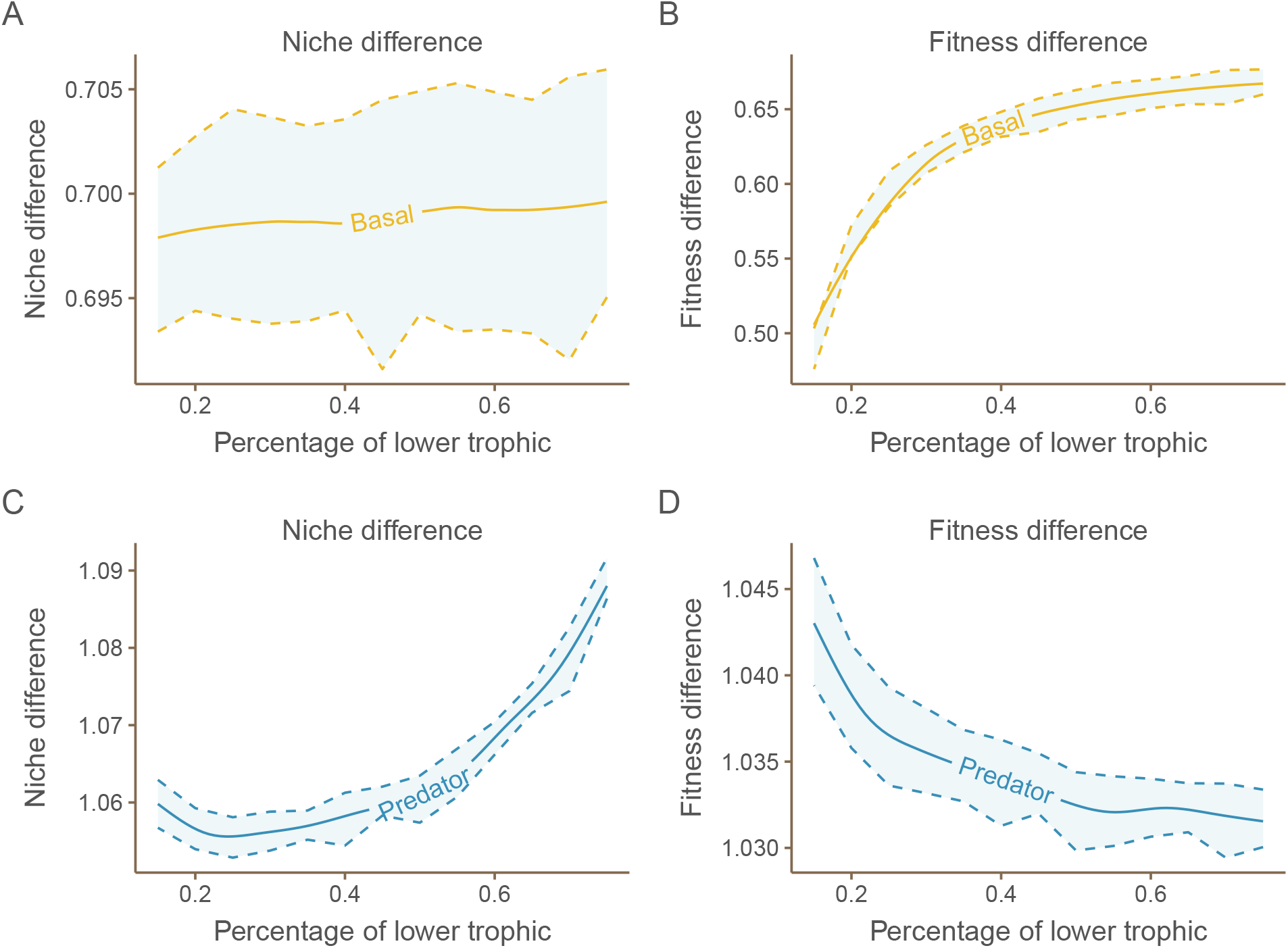
How community composition affects niche and fitness differences in two-trophic communities. We alter the trophic composition of the community while keeping total species richness constant. As with changes in species richness, the community composition does not affect niche differences of the lower trophic level (Panel A), but increases its fitness differences (Panel B). In contrast, niche differences of the higher trophic level increase with increasing proportion of basal species (Panel C), while fitness differences of the higher trophic level decrease with increasing proportion of basal species (Panel D). The pattern in Panel C is driven by the decrease of predator species richness (see Fig. 2 I), while the pattern in Panel D is driven by the increase of basal species richness (see Fig. 2 F). As with changes of species richness, the changes of niche differences are stronger than changes of fitness differences for the higher trophic level.

Focusing on fitness differences, increasing the proportion of basal species increases the fitness differences of the lower trophic level and slightly decreases fitness differences of the higher trophic level. Heuristically, increasing the proportion of basal species increases the total species density, which decreases the no-niche growth rate *η_i_*, a main driver of fitness differences (Eqn. 3). For a basal species, higher total density implies more fierce competition, which increases fitness differences. In contrast, for a predator species, higher total density implies more potential prey, which decreases their fitness differences.

The results obtained from the community focus are consistent with the traditional and alternative focuses, which suggest that a high or low proportion of basal species can hinder the persistence of either basal species or predators. Thus, our results predict two-trophic food webs have balanced proportion of basal and predator species. Notably, while the predicted patterns are the same, only by considering the community focus can we recognize that different trophic levels are influenced by distinct coexistence mechanisms.

### Tri-trophic communities and empirical data

In general, the results from three-trophic communities align with those of two-trophic communities. In both cases, trophic levels are more stable when they have lower species richness (Fig. 4G-I). Niche differences in the first trophic level remain unaffected by the trophic composition (Fig. 4 A), while the second and third levels are affected (Fig. 4B and C). The fitness differences of the second and third trophic levels are also minimally impacted by the trophic composition, except for a slight increase in basal species fitness when there is a high percentage of them present (Fig. 4D). These observations can be explained heuristically using similar reasoning as in two-trophic communities.

**Figure 4:**
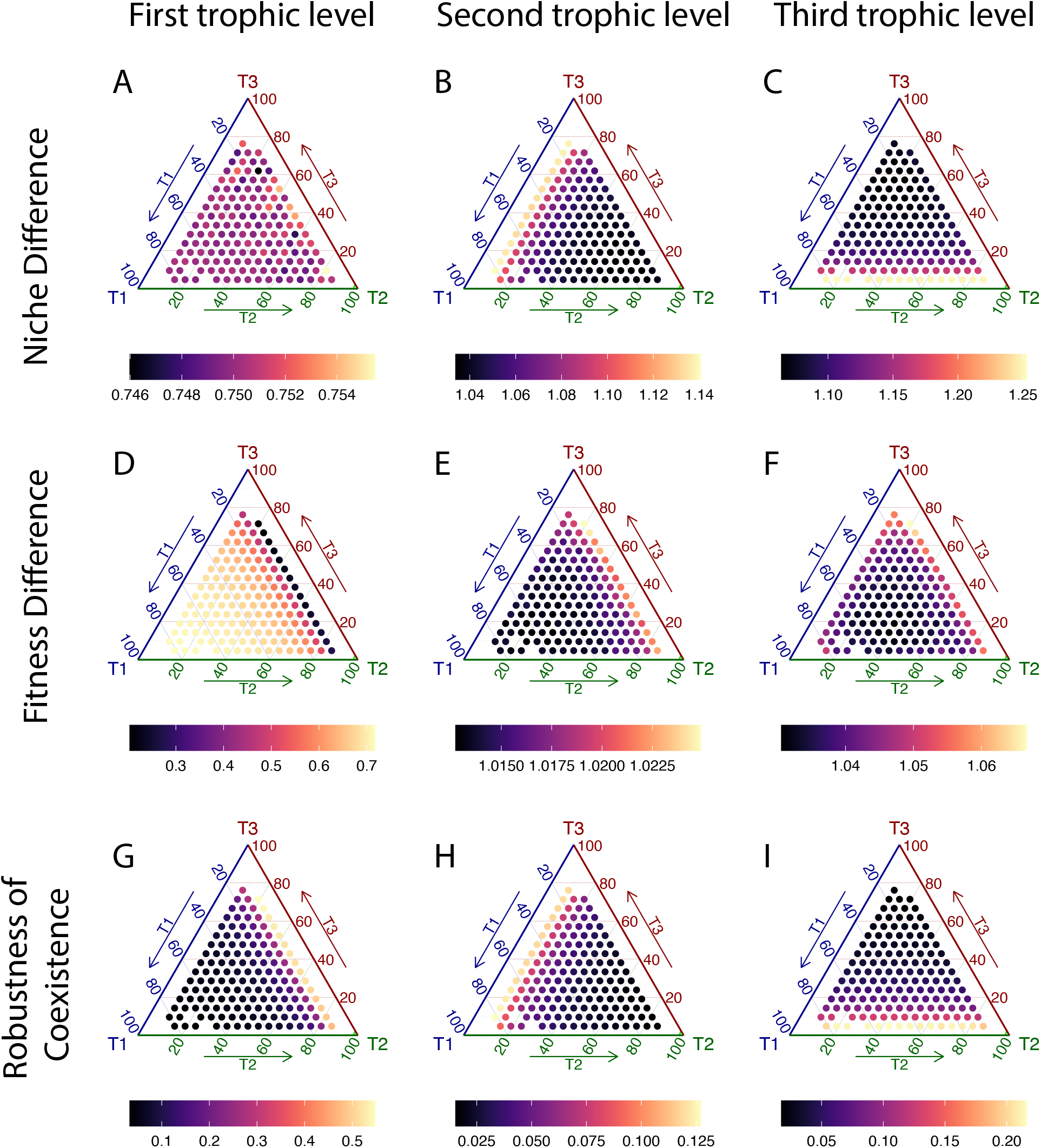
How community composition affects niche and fitness differences in three-trophic communities. We computed niche and fitness differences for communities with fixed number of species but varying tropical composition. The location in the ternary plots show the respective proportions of each trophic level. We computed niche (Panels A-C) and fitness differences (Panels D-F) for the species according to the community focus. As found with the two-trophic communities, community composition affects the fitness differences of the first trophic level (Panels D), but not the niche differences (Panels A). However, community composition affects niche differences and to a lesser extent fitness differences of the higher trophic levels (Panels B, C, E & F). Additionally, we also compute the stability of a species against extinction (Panels G-I), measured as how much niche difference overcompensates fitness difference 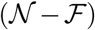. The color of the dots represent the stability of the least stable species per trophic level. As found in the two-trophic level communities, each trophic level is most stable if it contains relatively few species.

We analyzed 358 empirical predator-prey networks and found that they support the theoretical prediction of balanced species richness across trophic levels (Fig. 5). However, the empirical data suggests that the proportion of species in higher trophic levels is underestimated by the theoretical prediction of the most stable trophic composition (Fig. 5).

**Figure 5:**
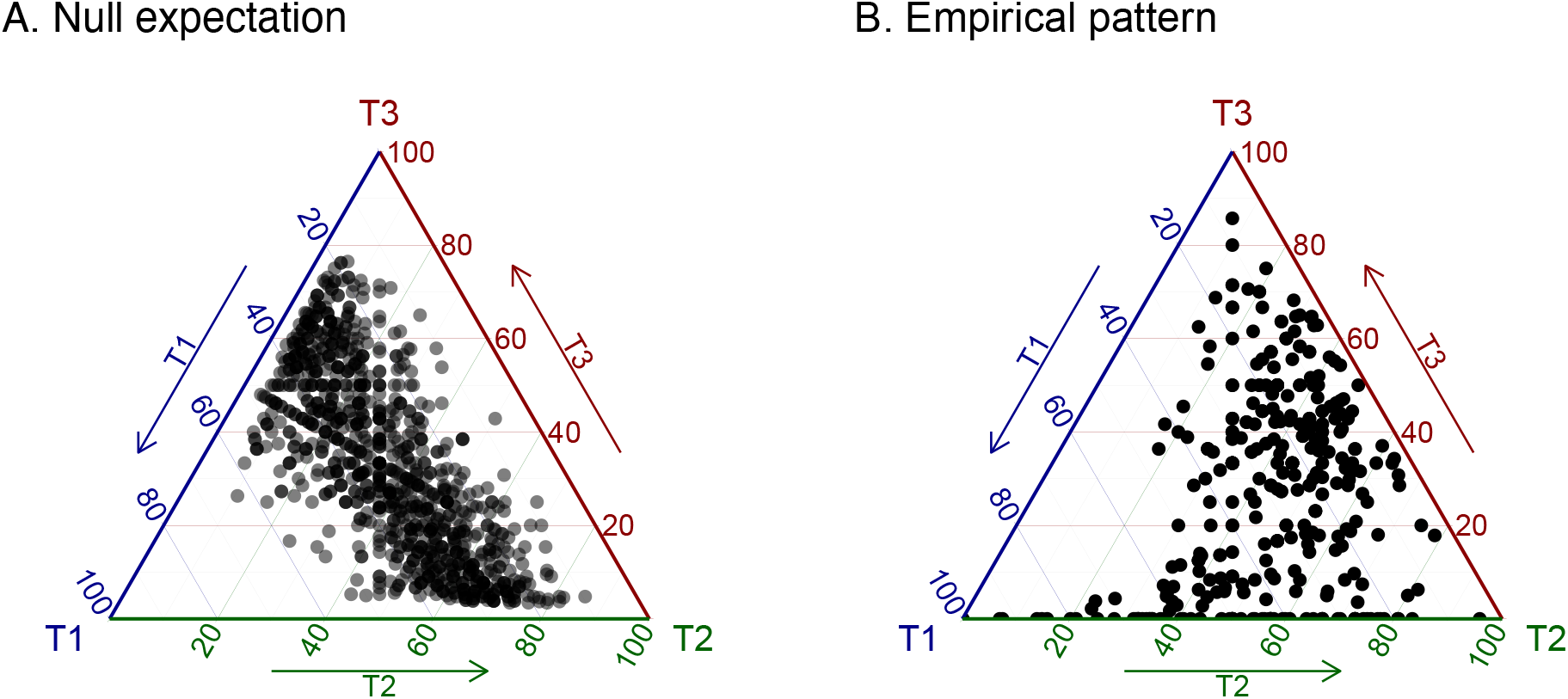
Comparing theoretical null expectation with empirical pattern. Panel A: For 500 different parameter settings we computed the top 5 most stable tropical configuration similar as in figure 4. Panel B: Tropical composition of empirical food webs from Web of science. Species with intermediate trophic levels were round to the nearest integer. Species with trophic levels 4 and higher were lumped with species from level 3. Color indicates species richness of the community, we excluded food-webs with less than 10 species.

## Discussion

We have filled a gap in modern coexistence theory on the null expectation of how assembly in one trophic level affects coexistence of other trophic levels or the community as a whole. We argue that studying the entire multitrophic structure is essential for many research questions, as it provides a comprehensive understanding of community dynamics (Godoy *et al*., 2018; Pande & Shnerb, 2022). For example, our findings demonstrate that relying solely on traditional or alternative approaches would lead to incorrect conclusions about the mechanisms driving coexistence across different trophic levels.

Our findings show that increasing species richness within a trophic level decreases its coexistence due to increased fitness differences and unaffected niche differences. Similarly, increasing species richness in an adjacent trophic level reduces stability because niche differences decrease more than fitness differences (Fig. 2). This is consistent with previous theoretical studies arguing that increasing species richness decreases stability (Spaak *et al*., 2021b; Allesina & Tang, 2015). However, it differs from previous empirical findings that the inclusion of a predator only affects fitness differences of the basal species, not however their niche differences (Petry *et al*., 2018; Terry *et al*., 2021).

Our theoretical analysis predicts a balanced proportion of basal and higher trophic species in food webs. We are not the first to find this relationship. Other different theoretical lens, such as Allesina & Tang (2012) using asymptotic dynamical stability and Carpentier *et al*. (2021) using robustness after extinctions, have made similar predictions. However, a key difference is that our work further provides a decomposition of this pattern to different forces in the different trophic levels: The diversity in the lowest trophic level is limited by increasing fitness differences (Fig. 2 D, F, &L, and 4 D&G), while the diversity in the higher trophic levels is limited by decreasing niche differences (Fig. 2 B, C, & I, and 4 E, F, H, & I).

Unfortunately, the traditional formalism of modern coexistence theory does not consider multitrophic structures. As a result, empirical studies that fall under this theory rarely include multi-trophic structures explicitly (Buche *et al*., 2022). Consequently, it is unclear whether the mechanisms responsible for maintaining coexistence differ from one trophic level to another. Our study provides a baseline of expected coexistence mechanisms from a null model of ecological assembly. Deviations from these baselines can indicate more complex ecological structures and mechanisms in nature.

### Limitations and future work

#### Our results are based on three critical simplifications of natural communities

First, we have adopted a simple trophic network structure. We did not consider omnivory, as all species in our simulations belonged to a specific trophic level. Additionally, each species interacted with each species from the adjacent trophic levels as well as with all species in the same trophic level. Such a simple network is tractable for our questions and highlight the current practice of traditional and alternative focus. However, empirical networks are typically sparse and also contain a significant amount of omnivory, such that a two species could have predator-prey relations, competitive relation for common prey and apparent competition with a common predator (Pimm *et al*., 1991; Holt & Bonsall, 2017). It is possible to extend our analysis with simulations with more realistically generated food-webs (Williams & Martinez, 2000; Allesina & Pascual, 2008; McCann, 2011). A caveat in that case, though, is that trophic level is not clearly defined. We hypothesize that many species with identical trophical identity, i.e. same prey and same predators, will make a food-network less stable. Similarly, we hypothesize that composition will affect fitness differences of basal species, but niche differences of predatory species.

Second, we assumed that all species interactions, independent of trophic level, are equally strong. This assumption is clearly unrealistic. However, we lack a decent empirical understanding of trophic interaction strengthen, as measuring empirical non-trophic interactions is challenging. For example, Kawatsu *et al*. (2021) found that roughly a fourth of all species interactions were not driven by trophic interactions. Spaak *et al*. (2022a) found that predation was much more important than resource competition in simulated plankton communities, but they did not include any non-trophic interactions.

Conceptually, species interactions can be disentangled into interactions with the higher trophic level, the lower trophic level and non-trophic interactions (Kawatsu *et al*., 2021). Mathematically, the effective interactions within a trophic level (*A*^11′^) can be decomposed into these respective parts, i.e. *A*^11′^ = *A*^11^ + *A*^12^(*A*^22^)^−1^*A*^21^ + *A*^10^(*A*^00^)^−1^*A*^01^, where *A*^00^, *A*^11^ and *A*^22^ capture non-trophic interactions, *A*^12^ and *A*^21^ capture interactions with the higher trophic level, and *A*^01^ and *A*^10^ capture interactions with the lower trophic level. While all these mechanisms affect *A*^11′^ similarly, the relative importance of these mechanisms is less clear. For simplicity, we here assumed that *A*^00^, *A*^11^ and *A*^22^ are all identically distributed. Yet, if we assume that interactions within a certain trophic level are much stronger, we should expect a lower richness in this trophic level.

Third, our analysis is based on Lotka-Volterra model. This model only captures linear species interactions, despite that non-linear species interactions are ubiquitous and the potentials for higher order interactions (Mickalide & Kuehn, 2019; Kleinhesselink *et al*., 2022). Unfortunately, computing niche and fitness differences for non-linear species interactions is challenging for various reasons. On the one hand, with nonlinear functional responses or higher order interactions, a community can have multiple stable equilibrium (*Barabás et al*., 2018; AlAdwani & Saavedra, 2020). This renders invasion growth rate, consequently niche and fitness differences, ill-defined.

## Author contributions

J.W.S. conceived the study. J.W.S. and C.S. performed the study. J.W.S. and C.S. wrote the manuscript.

