## Supplemental material for "Multitrophic assembly: a perspective from modern coexistence theory"

#### Contents

**A Model assumption** **S2**

**B Traditional focus** **S3**

**C Alternative focus** **S6**

**D Community focus** **S8**

**E Summary of key results** **S12**

### A Model assumption

We here present the analytical computations of how changes in species richness in the different trophic levels affect niche and fitness differences according to the different foci. For the analytic computation of niche and fitness differences we assume that all inter-specific interactions are identical, i.e. $A_{ij}^{xy} = a$, as opposed to a random variable with mean $a$. Furthermore, lete $n_1$ be the species richness in the first trophic level and $n_2$ be the species richness in the second trophic level. To compute niche and fitness differences according to the community focus we first compute the equilibrium densities of the species. Because of symmetry, all basal species must have the same density at equilibrium, i.e. $N_i^1 = N_j^1$ and similar for the second trophic level.

$$\frac{1}{N_i^1}\frac{dN_i^1}{dt} = 0 = \mu_i^1 - \sum_j A_{ij}^{11} N_j^1 - \sum_k A_{ik}^{12} N_k^2 \tag{S1}$$

$$= \mu_i^1 - (1 + a(n_1 - 1))N_i^1 - an_2 N_k^2 \tag{S2}$$

$$\frac{1}{N_i^2}\frac{dN_i^2}{dt} = 0 = \mu_i^2 - \sum_j A_{ij}^{21} N_j^1 - \sum_k A_{ik}^{22} N_k^2 \tag{S3}$$

$$= \mu_i^2 + an_1 N_j^1 - (1 + a(n_2 - 1))N_k^2 \tag{S4}$$

Solving the equation system we get

$$\begin{pmatrix} N_i^1 \\ N_i^2 \end{pmatrix} = \begin{pmatrix} 1 + a(n_1 - 1) & an_2 \\ -an_1 & 1 + a(n_2 - 1) \end{pmatrix}^{-1} \begin{pmatrix} \mu_1 \\ \mu_2 \end{pmatrix} \tag{S5}$$

$$= \frac{1}{(1 + a(n_1 - 1))(1 + a(n_2 - 1)) + a^2 n_1 n_2} \begin{pmatrix} 1 + a(n_2 - 1) & -an_2 \\ an_1 & 1 + a(n_1 - 1) \end{pmatrix} \begin{pmatrix} \mu_1 \\ \mu_2 \end{pmatrix} \tag{S6}$$

$$= \frac{1}{1 + a(n_1 - 1) + a(n_2 - 1) + a^2(n_1 - 1)(n_2 - 1) + a^2 n_1 n_2} \begin{pmatrix} an_2(\mu^1 - \mu^2) + (1 - a)\mu^1 \\ an_1(\mu^1 + \mu^2) + (1 - a)\mu^2 \end{pmatrix} \tag{S7}$$

$$= \frac{1}{(1 - a)^2 + a(1 - a)(n_1 + n_2) + 2a^2 n_1 n_2} \begin{pmatrix} an_2(\mu^1 - \mu^2) + (1 - a)\mu^1 \\ an_1(\mu^1 + \mu^2) + (1 - a)\mu^2 \end{pmatrix} \tag{S8}$$

We note that $N_i^2 > 0$ if and only if $n_1 > \frac{(1-a)|\mu_i^2|}{a(\mu_i^1 + \mu_i^2}$.

For the traditional and the alternative approach we first convert the community model with two trophic levels into a community model with only one trophic level by setting the growth rates of the other community level to constant. Often this is done with a timescale separation, which in this case would not be meaningful, as we cannot assume at the same time that the dynamics of the higher trophic level are faster as the basal and vice versa. Rather, the justification for this stems from the fact how we compute niche and fitness differences, which depends on the intrinsic growth rate $\mu_i$, the invasion growth rate $r_i$ and the no-niche growth rate $\eta_i$, which are all evaluated at an equilibrium state. We therefore do not assume timescale separation, but rather we only evaluate the growth dynamics at equilibrium, such that we can assume the non-focal trophic level to be at equilibrium.

### B Traditional focus

We first investigate how a change in richness of the lower trophic level affects niche and fitness differences of the lower trophic level. The density of the higher trophic level is given by

$$\frac{1}{N^2}\frac{dN^2}{dt} = \mu^2 - A^{21}N^1 - A^{22}N^2 = 0 \tag{S9}$$

$$\Leftrightarrow N^2 = (A^{22})^{-1}(\mu^2 - A^{21}N^1) \tag{S10}$$

Inserting this into the equation for the of the lower trophic level we get

$$\frac{1}{N^1}\frac{dN^1}{dt} = \mu^1 - A^{12}N^2 - A^{11}N^1 \tag{S11}$$

$$= \mu^1 - A^{11}N^1 - A^{12}(A^{22})^{-1}\mu^2 + A^{12}(A^{22})^{-1}A^{21}N^1 \tag{S12}$$

$$= (\mu^1 - A^{12}(A^{22})^{-1}\mu^2) - (A^{11} - A^{12}(A^{22})^{-1}A^{21})N^1 \tag{S13}$$

We therefore obtain another Lotka-Volterra community model with different parameters of the form ${\mu^1}' = \mu^1 - A^{12}(A^{22})^{-1}\mu^2$ and ${A^{11}}' = A^{11} - A^{12}(A^{22})^{-1}A^{21}$. To understand how species richness $n_1$ affects niche and fitness differences according to the traditional focus we must first understand how species richness $n_1$ affects ${A^{11}}'$.

We now compute ${A^{11}}' = A^{11} - A^{12}(A^{22})^{-1}A^{21}$. To do so, we note that $A^{22} = (1-a)I + aJ$, where $I$ is the unit matrix and $J$ is the matrix with all ones. We have $(A^{22})^{-1} = \frac{1}{1-a}\left(I - \frac{a}{1+(n_2-1)a}J\right)$, which can be verified as

$$A^{22}(A^{22})^{-1} = ((1-a)I + aJ)\frac{1}{1-a}\left(I - \frac{a}{1+(n_2-1)a}J\right) \tag{S14}$$

$$= I + \frac{1}{1-a}\left(-\frac{(1-a)a}{1+(n_2-1)a} + a - \frac{n_2a^2}{1+(n_2-1)a}\right) \tag{S15}$$

$$= I + \frac{1}{1-a}\left(\frac{-a + a^2 + a + (n_2-1)a^2 - n_2a^2}{1+(n_2-1)a}\right) \tag{S16}$$

$$= I \tag{S17}$$

Given this, we can compute $A^{12}(A^{22})^{-1}A^{21}$ as

$$A^{12}(A^{22})^{-1}A^{21} = A^{12}\frac{1}{1-a}\left(I - \frac{a}{1+(n_2-1)a}J\right)A^{21} \tag{S18}$$

$$= aJ\frac{1}{1-a}\left(I - \frac{a}{1+(n_2-1)a}J\right)(-aJ) \tag{S19}$$

$$= \frac{-a^2}{1-a}\left(JIJ - \frac{a}{1+(n_2-1)a}J\cdot J\cdot J\right) \tag{S20}$$

$$= \frac{-a^2}{1-a}\left(n_2 J - \frac{n_2 a}{1+(n_2-1)a}n_2^2 J\right) \tag{S21}$$

$$= -\frac{a^2}{1-a}\left(n_2 - \frac{n_2 a}{1+(n_2-1)a}\right)J \tag{S22}$$

$$= -\frac{a^2}{1-a}\frac{n_2 + n_2^2 a - n_2 a + n_2^2 a}{1+(n2-1)a}J \tag{S23}$$

$$= \frac{-n_2 a^2}{1+(n_2-1)a}J \tag{S24}$$

Given this we can compute $A^{11'}_{ij} = a + \frac{n_2 a^2}{1+(n_2-1)a}$ and $A^{11'}_{ii} = 1 + \frac{n_2 a^2}{1+(n_2-1)a}$.

We can then compute niche differences of the first trophic level as

$$\mathcal{N}_i^1 = \frac{r_i - \eta_i}{\mu_i - \eta_i} \tag{S25}$$

$$= \frac{(\mu_i' - n_1 A^{11'}_{ij} N^1) - (\mu_i' - n_1 A^{11'}_{ii} N^1)}{(\mu_i') - (\mu_i' - n_1 A^{11'}_{ii} N^1)} \tag{S26}$$

$$= 1 - \frac{A^{11'}_{ij}}{A^{11'}_{ii}} \tag{S27}$$

We see that this does not depend on $n_1$, the species richness in the first trophic level, as neither $A^{11'}$ is independent of $n_1$.

To compute $\mathcal{F}_i^2$ we first compute the equilibrium density of the species as

$$0 = \mu^{1'} - A^{11'} N^{1'} \tag{S28}$$

$$= \mu^{2'} - A^{11'}_{ij}(n_1 - 1)N^{1'} - A^{11'}_{ii} N^{1'} \tag{S29}$$

$$N^{1'} = \frac{\mu_i^{1'}}{A^{11'}_{ii} + (n_1-1)A^{11'}_{ij}} \tag{S30}$$

we therefore get $\eta_i = \mu_i^{1'} - \frac{A^{11'}_{ii} n_1 \mu_i^{1'}}{A^{11'}_{ii} + (n_1-1)A^{11'}_{ij}}$

$$\mathcal{F}_i^1 = 1 - \frac{\mu_i}{\mu_i - \eta_i} \tag{S31}$$

$$= 1 - \frac{\mu_i'}{\frac{A_{ii}^{11'} n_1 \mu_i^{1'}}{A_{ii}^{11'} + (n_1 - 1) A_{ij}^{11'}}} \tag{S32}$$

$$= 1 - \frac{1}{n_1} - \frac{(n_1 - 1) A_{ij}^{11'}}{n_1 A_{ii}^{11'}} \tag{S33}$$

$$= \left(1 - \frac{A_{ij}^{11'}}{A_{ii}^{11'}}\right)\left(1 - \frac{1}{n_1}\right) \tag{S34}$$

We therefore see that increasing $n_1$ increases $\mathcal{F}_i^1$. The same analysis applies if we change the species richness of the higher trophic level and analyse the niche and fitness differences of the higher trophic level. For the traditional focus we therefore see that the results from Spaak *et al.* (2021) apply, that is species richness in the same trophic level have no effect on niche differences, but does increase fitness differences.

### C Alternative focus

Similar to the traditional focus we can compute $A^{22'} = A^{22} - A^{21}(A^{11})^{-1}A^{12}$ which gives

$$A^{22'}_{ij} = a + \frac{n_1 a^2}{1 + (n_1 - 1)a} \tag{S35}$$

$$= \frac{a + n_1 a^2 - a^2 + n_1 a^2}{1 + (n_1 - 1)a} \tag{S36}$$

$$= a\frac{1 - a + 2n_1 a}{1 + (n_1 - 1)a} \tag{S37}$$

$$A^{22'}_{ii} = 1 + \frac{n_1 a^2}{1 + (n_1 - 1)a} \tag{S38}$$

$$= \frac{1 + (n_1 - 1)a + n_1 a^2}{1 + (n_1 - 1)a} \tag{S39}$$

$$= \frac{1 - a + a n_1(1 + a)}{1 + (n_1 - 1)a} \tag{S40}$$

Next we can compute the relative interspecific competition, i.e.

$$\frac{A^{22'}_{ij}}{A^{22'}_{ii}} = \frac{a\,(1 - a + 2n_1 a)}{1 - a + a(1 + a)n_1} \tag{S41}$$

$$= a\frac{1 - a + 2n_1 a + a(1 + a)n_1 - a(1 + a)n_1}{1 - a + a(1 + a)n_1} \tag{S42}$$

$$= a\left(1 + \frac{2n_1 a - a(1 + a)n_1}{1 - a + a(1 + a)n_1}\right) \tag{S43}$$

$$= a\left(1 + \frac{a(1 - a)n_1}{1 - a + a(1 + a)n_1}\right) \tag{S44}$$

Therefore, the relative interspecific competition strength is increasing in the species richness of the lower trophic level. As a consequence, we expect niche differences to decrease with increasing $n_1$.

To compute $\mathcal{F}^2_i$ we note that $c_{ij} = 1$ because of symmetry. To compute the no-niche growth rate we first compute the equilibrium density of the species as

$$0 = \mu^{2'} - A^{22'}N^{2'} \tag{S45}$$

$$= \mu^{2'} - A^{22'}_{ij}(n_2 - 1)N^{2'} - A^{22'}_{ii}N^{2'} \tag{S46}$$

$$N^{2'} = \frac{\mu^{2'}}{A^{22'}_{ii} + (n_2 - 1)A^{22'}_{ij}} \tag{S47}$$

We therefore get $\eta_i = \mu^{2'} - \frac{A^{22'}_{ii} n_2 \mu^{2'}}{A^{22'}_{ii} + (n_2 - 1)A^{22'}_{ij}}$ and therefore

$$\mathcal{F}_i^2 = 1 - \frac{\mu_i}{\mu_i - \eta_i} \tag{S48}$$

$$= 1 - \frac{\mu^{2'}}{\frac{n_2 \mu^{2'} A_{ii}^{22'}}{A_{ii}^{22'} + (n_2 - 1) A_{ij}^{22'}}} \tag{S49}$$

$$= 1 - \frac{1 + (n_2 - 1)\frac{A_{ij}^{22'}}{A_{ii}^{22'}}}{n_2} \tag{S50}$$

$$= 1 - \frac{A_{ij}^{22'}}{A_{ii}^{22'}} - \frac{1 - \frac{A_{ij}^{22'}}{A_{ii}^{22'}}}{n_2} \tag{S51}$$

$$= \left(1 - \frac{A_{ij}^{22'}}{A_{ii}^{22'}}\right)\left(1 - \frac{1}{n_2}\right) \tag{S52}$$

As shown above, $1 - \frac{A_{ij}^{22'}}{A_{ii}^{22'}}$ decreases in $n_1$, therefore fitness differences decrease with increasing species richness of the adjacent trophic level.

Conversely, focusing on the effect of changing the species richness in the higher trophic level and assessing it's effect on the lower trophic level. We know that $\frac{A_{ij}^{11'}}{A_{ii}^{11'}}$ increases in $n_2$, the species richness in the higher trophic level. Therefore, we expect niche differences of the lower trophic level to decrease with increasing species richness of the higher trophic level. For the same arguments as mentioned above we therefore expect fitness differences to decrease as well.

#### D Community focus

We note that the $c_{ij} = 1$ for all species, as all inter-specific species interactions have the same strength. With this we can compute the relevant niche and fitness differences of the two trophic levels

$$\mathcal{N}_i^1 = 1 - \frac{\mu_i^1 - r_i^1}{\mu_i^1 - \eta_i^1} \tag{S53}$$

$$= 1 - \frac{\mu_i^1 - (\mu_i^1 - an_1N^1 - an_2N^2)}{\mu_i^1 - (\mu_i^1 - n_1N^1 - n_2N^2)} \tag{S54}$$

$$= 1 - a \tag{S55}$$

Which is independent of species richness $n_1$ and $n_2$.

For the other niche and fitness differences we first observe that the derivative of the function $\frac{d}{dx}\left(-\frac{\alpha x+\beta}{\gamma x+\delta}\right) = -\frac{\alpha(\gamma x+\delta)-(\alpha x+\beta)\gamma}{(\gamma x+\delta)^2} = -\frac{\alpha\delta-\beta\gamma}{(\gamma x+\delta)^2}$ is negative if $\alpha\delta - \beta\gamma > 0$. Therefore, $-\frac{\alpha x+\beta}{\gamma x+\delta}$ is a decreasing function if $\alpha\delta - \beta\gamma > 0$.

$$\mathcal{F}_i^1 = 1 - \frac{\mu_i^1}{\mu_i^1 - \eta_i^1} \tag{S56}$$

$$= 1 - \frac{\mu_i^1}{\mu_i^1 - (\mu_i^1 - n_1N^1 - n_2N^2)} \tag{S57}$$

$$= 1 - \frac{\mu_i^1}{n_1N_i^1 + n_2N_i^2} \tag{S58}$$

$$= 1 - \mu_i^1 \frac{(1-a)^2 + a(1-a)(n_1+n_2) + 2a^2n_1n_2}{2an_1n_2\mu_i^1 + (1-a)(n_1\mu_i^1 + n_2\mu_i^2)} \tag{S59}$$

$$= 1 - \mu_i^1 \frac{(1-a)^2 + \frac{a}{\mu_i^1}\left((1-a)(n_1+n_2)\mu_i^1 + 2an_1n_2\mu_i^1\right)}{2an_1n_2\mu_i^1 + (1-a)(n_1\mu_i^1 + n_2\mu_i^2)} \tag{S60}$$

$$= 1 - \mu_i^1 \left(\frac{a}{\mu_i^1} + \frac{(1-a)^2 + \frac{a}{\mu_i^1}\left((1-a)n_2\mu_i^1 - (1-a)n_2\mu_i^2\right)}{2an_1n_2\mu_i^1 + (1-a)(n_1\mu_i^1 + n_2\mu_i^2)}\right) \tag{S61}$$

$$= 1 - a - \frac{\mu_i^1(1-a)^2 + a(1-a)n_2(\mu_i^1 - \mu_i^2)}{2an_1n_2\mu_i^1 + (1-a)(n_1\mu_i^1 + n_2\mu_i^2)} \tag{S62}$$

$$= (1-a)\left(1 - \frac{\mu_i^1(1-a) + an_2(\mu_i^1 - \mu_i^2)}{2an_1n_2\mu_i^1 + (1-a)(n_1\mu_i^1 + n_2\mu_i^2)}\right) \tag{S63}$$

We therefore see that increasing $n_1$ increases $\mathcal{F}_i^1$, as the last term in the brackets is of form $-\frac{0n_1+\beta}{\gamma n_1+\delta}$ with $\beta, \gamma, \delta > 0$. Additionally, for increasing $n_1$ $\mathcal{F}_i^1$ approaches $1-a$. To assess the effect of $n_2$ on $\mathcal{F}_i^1$ we note that the last term has the form $-\frac{\alpha n_2+\beta}{\gamma n_2+\delta}$ with $\alpha = a(\mu_i^1 - \mu_i^2), \beta = \mu_i^1(1-a)$,

$\gamma = 2an_1\mu_i^1 + (1-a)\mu_i^2, \delta = (1-a)n_1\mu_i^1$, which leads to

$$\alpha\delta - \beta\gamma = a(\mu_i^1 - \mu_i^2)(1-a)n_1\mu_i^1 - \mu_i^1(1-a)(2an_1\mu_i^1 + (1-a)\mu_i^2) \tag{S64}$$
$$= (1-a)\mu_i^1\left(an_1(\mu_i^1 - \mu_i^2 - 2\mu_i^1) - (1-a)\mu_i^2\right) \tag{S65}$$
$$= (1-a)\mu_i^1\left(-an_1(\mu_i^1 + \mu_i^2) - (1-a)\mu_i^2\right) \tag{S66}$$

This is negative if $n_1 > \frac{(1-a)|\mu_i^2|}{a(\mu_i^1+\mu_i^2)}$. This condition is equivalent to the condition that the higher trophic level have a positive equilibrium density, i.e. $N_i^2 > 0$, see S8 We therefore conclude that $F_i^1$ increases with increasing $n_2$.

$$\mathcal{N}_i^2 = 1 - \frac{\mu_i^2 - r_i^2}{\mu_i^2 - \eta_i^2} \tag{S67}$$
$$= 1 - \frac{-an_1N_i^1 + an_2N_i^2}{n_1N_i^1 + n_2N_i^2} \tag{S68}$$
$$= 1 - a\frac{-n_1N_i^1 + n_2N_i^2}{n_1N_i^1 + n_2N_i^2} \tag{S69}$$
$$= 1 - a\frac{-n_1(an_2(\mu_i^1 - \mu_i^2) + (1-a)\mu_i^1) + n_2(an_1(\mu_i^1 + \mu_i^2) + (1-a)\mu_i^2)}{n_1(an_2(\mu_i^1 - \mu_i^2) + (1-a)\mu_i^1) + n_2(an_1(\mu_i^1 + \mu_i^2) + (1-a)\mu_i^2)} \tag{S70}$$
$$= 1 - a\frac{2\mu_i^2an_1n_2 - (1-a)(n_1\mu_i^1 - n_2\mu_i^2)}{2\mu_i^1an_1n_2 + (1-a)(n_1\mu_i^1 + n_2\mu_i^2)} \tag{S71}$$
$$= 1 - a\left(\frac{\mu_i^2}{\mu_i^1} + \frac{-(1-a)(n_1\mu_i^1 - n_2\mu_i^2) - (1-a)(n_1\mu_i^1 + n_2\mu_i^2)\frac{\mu_i^2}{\mu_i^1}}{2\mu_i^1an_1n_2 + (1-a)(n_1\mu_i^1 + n_2\mu_i^2)}\right) \tag{S72}$$
$$= 1 - a\left(\frac{\mu_i^2}{\mu_i^1} - (1-a)\frac{n_1(\mu_i^1 + \mu_i^2) - n_2\mu_i^2\frac{\mu_i^1 - \mu_i^2}{\mu_i^1}}{2\mu_i^1an_1n_2 + (1-a)(n_1\mu_i^1 + n_2\mu_i^2)}\right) \tag{S73}$$
$$= 1 - a\frac{\mu_i^2}{\mu_i^1} - \frac{a(1-a)n_2\mu_i^2\frac{\mu_i^1 - \mu_i^2}{\mu_i^1}}{2\mu_i^1an_1n_2 + (1-a)(n_1\mu_i^1 + n_2\mu_i^2)} + \frac{a(1-a)n_1(\mu_i^1 + \mu_i^2)}{2\mu_i^1an_1n_2 + (1-a)(n_1\mu_i^1 + n_2\mu_i^2)} \tag{S74}$$
$$= 1 - a\frac{\mu_i^2}{\mu_i^1} - a(1-a)\frac{n_2\mu_i^2\frac{\mu_i^1 - \mu_i^2}{\mu_i^1} - n_1(\mu_i^1 + \mu_i^2)}{2\mu_i^1an_1n_2 + (1-a)(n_1\mu_i^1 + n_2\mu_i^2)} \tag{S75}$$

The last expression is of the form $-\frac{\alpha n_1+\beta}{\gamma n_1+\delta}$ with $\alpha = -(\mu_i^1 + \mu_i^2), \beta = n_2\mu_i^2\frac{\mu_i^1-\mu_i^2}{\mu_i^1}$, $\gamma = 2a\mu_i^1n_2 + (1-a)\mu_i^1, \delta = (1-a)n_2\mu_i^2$ where $\alpha, \beta$ and $\delta$ are negative, hence $\alpha\delta - \beta\gamma$ is positive and hence $\mathcal{N}_i^2$ decreases with increasing $n_1$.

To assess the dependence on $n_2$ we not that the last expression is of the form $-\frac{\alpha n_2+\beta}{\gamma n_2+\delta}$ with

$\alpha = \mu_i^2 \frac{\mu_i^1 - \mu_i^2}{\mu_i^1}, \beta = -n_1(\mu_i^1 + \mu_i^2)$, $\gamma = 2\mu_i^1 a n_1 + (1-a)\mu_i^2, \delta = (1-a)n_1\mu_i^1$ which leads to

$$\alpha\delta - \beta\gamma = \mu_i^2 \frac{\mu_i^1 - \mu_i^2}{\mu_i^1}(1-a)n_1\mu_i^1 - (-n_1(\mu_i^1 + \mu_i^2))(2\mu_i^1 a n_1 + (1-a)\mu_i^2) \tag{S76}$$

$$= n_1 \left((1-a)\mu_i^2((\mu_i^1 - \mu_i^2) + (\mu_i^1 + \mu_i^2)) + (\mu_i^1 + \mu_i^2)2\mu_i^1 a n_1\right) \tag{S77}$$

$$= 2n_1\mu_i^1 \left((1-a)\mu_i^2 + a n_1(\mu_i^1 + \mu_i^2)\right) \tag{S78}$$

This last expression is positive if $n_1 > \frac{|\mu_i^2|(1-a)}{a(\mu_i^1 + \mu_i^2)}$, which is the same expression as for $\mathcal{F}_i^1$. We therefore note that $\mathcal{N}_i^2$ decreases in $n_2$.

$$\mathcal{F}_i^2 = 1 - \frac{\mu_i^2}{\mu_i^2 - \eta_i^2} \tag{S79}$$

$$= 1 - \frac{\mu_i^2}{n_1 N_i^1 + n_2 N_i^2} \tag{S80}$$

$$= 1 - \mu_i^2 \frac{(1-a)^2 + a(1-a)(n_1 + n_2) + 2a^2 n_1 n_2}{2a n_1 n_2 \mu_i^1 + (1-a)(n_1\mu_i^1 + n_2\mu_i^2)} \tag{S81}$$

$$= 1 - \mu_i^2 \frac{(1-a)^2 + \frac{a}{\mu_i^1}\left((1-a)(n_1 + n_2)\mu_i^1 + 2a n_1 n_2 \mu_i^1\right)}{2a n_1 n_2 \mu_i^1 + (1-a)(n_1\mu_i^1 + n_2\mu_i^2)} \tag{S82}$$

$$= 1 - \mu_i^2 \left(\frac{a}{\mu_i^1} + \frac{(1-a)^2 + \frac{a}{\mu_i^1}\left((1-a)n_2\mu_i^1 - (1-a)n_2\mu_i^2\right)}{2a n_1 n_2 \mu_i^1 + (1-a)(n_1\mu_i^1 + n_2\mu_i^2)}\right) \tag{S83}$$

$$= 1 - a\frac{\mu_i^2}{\mu_i^1} - (1-a)\frac{\mu_i^2(1-a) + \mu_i^2 a n_2 \frac{\mu_i^1 - \mu_i^2}{\mu_i^1}}{2a n_1 n_2 \mu_i^1 + (1-a)(n_1\mu_i^1 + n_2\mu_i^2)} \tag{S84}$$

$$= 1 - a\frac{\mu_i^2}{\mu_i^1} - \frac{a(1-a)n_2\mu_i^2 \frac{\mu_i^1 - \mu_i^2}{\mu_i^1}}{2\mu_i^1 a n_1 n_2 + (1-a)(n_1\mu_i^1 + n_2\mu_i^2)} - \frac{\mu_i^2(1-a)^2}{2\mu_i^1 a n_1 n_2 + (1-a)(n_1\mu_i^1 + n_2\mu_i^2)} \tag{S85}$$

$$\tag{S86}$$

The expression in S84is of the form $-\frac{0n_1 + \beta}{\gamma n_1 + \delta}$ with negative $\beta$ (note that $\mu_i^2 < 0$) and positive $\gamma$, therefore $\mathcal{F}_i^2$ decreases in $n_1$.

The expression in S84is of the form $\frac{\alpha n_2 + \beta}{\gamma n_1 + \delta}$ with $\alpha = \mu_i^2 a \frac{\mu_i^1 - \mu_i^2}{\mu_i^1}, \beta = \mu_i^2(1-a)$, $\gamma = 2a n_1 \mu_i^1 + (1-a)\mu_i^2$ and $\delta = (1-a)n_1\mu_i^1$ which leads to

$$\alpha\delta - \beta\gamma = \mu_i^2 a \frac{\mu_i^1 - \mu_i^2}{\mu_i^1}(1-a)n_1\mu_i^1 - \mu_i^2(1-a)(2a n_1\mu_i^1 + (1-a)\mu_i^2) \tag{S87}$$

$$= (1-a)\mu_i^2 \left(a n_1(\mu_i^1 - \mu_i^2) - 2a n_1\mu_i^1 - (1-a)\mu_i^2\right) \tag{S88}$$

$$= -(1-a)\mu_i^2 \left(a n_1(\mu_i + \mu_i^2) + (1-a)\mu_i^2\right) \tag{S89}$$

This last expression is positive if $n_1 > \frac{|\mu_i^2|(1-a)}{a(\mu_i^1 + \mu_i^2)}$, which is the same expression as for $\mathcal{F}_i^1$ and for $\mathcal{N}_i^2$. We therefore note that $\mathcal{N}_i^2$ decreases in $n_2$.

##### D.1 Relative change of niche and fitness differences

For the higher trophic level we note that the expression for niche and fitness differences are almost equivalent (see equations S74 and S85). Specifically, they differ only in the last term of each expression.

$$\mathcal{F}_i^2 - \mathcal{N}_i^2 = -\frac{a(1-a)n_1(\mu_i^1 + \mu_i^2)}{2\mu_i^1 a n_1 n_2 + (1-a)(n_1\mu_i^1 + n_2\mu_i^2)} - \frac{\mu_i^2(1-a)^2}{2\mu_i^1 a n_1 n_2 + (1-a)(n_1\mu_i^1 + n_2\mu_i^2)} \tag{S90}$$

$$= -\frac{(1-a)\left(an_1(\mu_i^1 + \mu_i^2) + \mu_i^2(1-a)\right)}{2\mu_i^1 a n_1 n_2 + (1-a)(n_1\mu_i^1 + n_2\mu_i^2)} \tag{S91}$$

Which is negative if $n_1 > \frac{|\mu_i^2|(1-a)}{a(\mu_i^1+\mu_i^2)}$, i.e. if the higher trophic level has positive equilibrium density. This expression is of the form $-\frac{\alpha n_1+\beta}{\gamma n_1+\delta}$ with negative $\alpha, \gamma$ and $\delta$, but positive $\beta$, hence $\alpha\delta - \gamma\beta > 0$. Consequentially, $\frac{d}{dn_1}\mathcal{F}_i^2 - \frac{d}{dn_1}\mathcal{N}_i^2$ is negative, or said differently, changes in species richness of the lower trophic level have a stronger effect on niche than on fitness differences of the higher trophic level.

Similarly, this expression is of the form $-\frac{0n_2+\beta}{\gamma n_2+\delta}$ with positive $\beta$ and $\gamma$, hence $\alpha\delta - \gamma\beta < 0$. Consequentially, $\frac{d}{dn_1}\mathcal{F}_i^2 - \frac{d}{dn_1}\mathcal{N}_i^2$ is positive, or said differently, changes in species richness of the higher trophic level have a stronger effect on fitness than on niche differences of the higher trophic level.

### E Summary of key results

| Method | type | Equation | Effect low | Effect high |
|---|---|---|---|---|
| Trad. | $\mathcal{N}_i^1$ | $1-\frac{A_{ij}^{11\prime}}{A_{ii}^{11\prime}}$ | Constant | NA |
| Trad. | $\mathcal{F}_i^1$ | $\left(1-\frac{A_{ij}^{11\prime}}{A_{ii}^{11\prime}}\right)\left(1-\frac{1}{n_1}\right)$ | Increase | NA |
| Trad. | $\mathcal{N}_i^2$ | $1-\frac{A_{ij}^{22\prime}}{A_{ii}^{22\prime}}$ | NA | Constant |
| Trad. | $\mathcal{F}_i^2$ | $\left(1-\frac{A_{ij}^{22\prime}}{A_{ii}^{22\prime}}\right)\left(1-\frac{1}{n_2}\right)$ | NA | increase |
| Alt. | $\mathcal{N}_i^1$ | $1-a\left(\frac{a(1-a)n_2}{1-a+a(1+a)n_2}\right)$ | NA | decrease |
| Alt. | $\mathcal{F}_i^1$ | $\left(1-a\left(\frac{a(1-a)n_2}{1-a+a(1+a)n_2}\right)\right)\left(1-\frac{1}{n_1}\right)$ | NA | decrease |
| Alt. | $\mathcal{N}_i^2$ | $1-a\left(\frac{a(1-a)n_1}{1-a+a(1+a)n_1}\right)$ | decrease | NA |
| Alt. | $\mathcal{F}_i^2$ | $\left(1-a\left(\frac{a(1-a)n_1}{1-a+a(1+a)n_1}\right)\right)\left(1-\frac{1}{n_2}\right)$ | decrease | NA |
| Com. | $\mathcal{N}_i^1$ | $1-a$ | Constant | Constant |
| Com. | $\mathcal{F}_i^1$ | $(1-a)\left(1-\frac{\mu_i^1(1-a)+an_2(\mu_i^1-\mu_i^2)}{2an_1n_2\mu_i^1+(1-a)(n_1\mu_i^1+n_2\mu_i^2)}\right)$ | increase | increase |
| Com. | $\mathcal{N}_i^2$ | $1-a\frac{\mu_i^2}{\mu_i^1}-a(1-a)\frac{n_2\mu_i^2\frac{\mu_i^1-\mu_i^2}{\mu_i^1}-n_1(\mu_i^1+\mu_i^2)}{2\mu_i^1an_1n_2+(1-a)(n_1\mu_i^1+n_2\mu_i^2)}$ | decrease | decrease |
| Com. | $FD_i^2$ | $1-a\frac{\mu_i^2}{\mu_i^1}-(1-a)\frac{\mu_i^2(1-a)+\mu_i^2an_2\frac{\mu_i^1-\mu_i^2}{\mu_i^1}}{2an_1n_2\mu_i^1+(1-a)(n_1\mu_i^1+n_2\mu_i^2)}$ | decrease | decrease |

Table S1: We summarize the expressions for niche and fitness differences (second and third column) according to the three different focus (first column). We report the effect of increasing the lower (”Effect low”) or increasing the higher (”Effect high”) trophic level. NA implies that this focus can not be applied to the given change in species richness. For derivations see the text.
